# Comparative Analysis of Neural Crest Development in the Chick and Mouse

**DOI:** 10.1101/2024.11.06.622355

**Authors:** JA Morrison, I Pushel, R McLennan, MC McKinney, MM Gogol, A Scott, R Krumlauf, PM Kulesa

## Abstract

A core framework of the gene regulatory network (GRN) governing neural crest (NC) cell development has been generated by integrating separate inputs from diverse model organisms rather than direct comparison. This has limited insights into the diversity of genes in the NC cell GRN and extent of conservation of newly identified transcriptional signatures in cell differentiation and invasion. Here, we address this by leveraging the strengths and accessibility of the avian embryo to precise developmental staging by egg incubation and use an integrated analysis of chick (HH13) and mouse (E9.5) embryo tissue samples collected during NC cell migration into pharyngeal arches 1-2 (PA1 and PA2). We successfully identify a cluster of NC cells containing both mouse and chick cells that share expression of *Lmo4*, *Tfap2B*, *Sox10*, and *Twist1*, and distinct genes that lack known conserved roles in NC. Importantly, we discovered a cluster of cells exhibiting a conserved transcriptional signature associated with the NC cell migratory wavefront in both mouse and chick, including KAZALD1, BAMBI, DES, and GPC3. We confirm their expression is restricted to leader mouse NCs by multiplexed FISH. Together, these data offer novel insights into the transcriptional programs that underlie NC cell migration and establish the foundation for future comparative functional analyses.

## INTRODUCTION

Experiments with a diverse array of vertebrate model organisms have revealed core gene regulatory networks (GRN) governing the progressive steps of neural crest (NC) cell development (Simoes-Costa et al., 2014; Lumb et al., 2017; Plouhinec et al., 2017; Williams et al., 2019; Soldatov et al., 2019; Etchevers et al., 2019; Lencer et al., 2021). Moreover, existing evidence shows there are differences in the roles of key genes and signals when comparing the mouse NC cell GRN with other commonly studied vertebrates, such as chick, zebrafish and Xenopus (Barriga et al., 2015; Zhao and Trainor, 2022). This highlights the challenges to extrapolating between species based on the broad and simple framework of the current core NC cell GRN, generated by analyses in diverse species. Thus, there is a critical need to directly compare data at analogous stages of NC cell development in closely stage-matched samples of two organisms to better understand the degree of conservation in the components, signaling inputs and regulatory steps that underlie specific processes in the hierarchy of the progressive GRN architecture.

One of the intriguing features of NC cell development that is shared between organisms is the sculpting and long-distance invasion and migration of cells that extend from the dorsal neural tube to peripheral targets all along the vertebrate axis. Mistakes in NC cell migration lie at the center of many human birth defects in heart, nervous system, skin pigmentation and craniofacial development (Vega-Lopez et al., 2018; Siismets and Hatch, 2020). The diverse molecular mechanisms governing the interplay of bi-directional signaling between the migrating NC cells and their microenvironment(s), which ensure cell invasion and collective migration to precise locations in the embryo, are continuing to emerge from single cell (sc) RNA-seq studies (Lumb et al., 2017; Plouhinec et al., 2017; Williams et al., 2019). However, a more complete understanding of the properties of migrating NC cells will require deeper analyses of the spatial and temporal patterns of gene expression within NC cell streams and comparative approaches across organisms with the potential to shed light on the mechanistic basis of this phenomenon and its degree of conservation.

We previously leveraged the accessibility and ease of tissue manipulation of the avian model organism to perform a scRNA-seq analysis of cranial NC cell migration at three progressive stages in chick (HH11, 13, 15; Hamburger and Hamilton, 1951) and established hierarchical relationships between cell position and time-specific transcriptional signatures (Morrison et al., 2017). These data revealed molecular diversity and dynamics within a NC cell migratory stream that underlie complex directed and collective cell behaviors, including a novel transcriptional signature of the most invasive (we termed Trailblazer NC cells) that is consistent during migration. This stimulated us to derive the transcriptional states of migrating NC cells and the cellular landscape of the first four chick cranial to cardiac branchial arches (BA1-4) using label free, unsorted scRNA sequencing (Morrison et al., 2021). The faithful capture of branchial arch specific genes led to the identification of novel markers of migrating NC cells and Trailblazer invasion genes common to all BA1-4 streams but did not include a direct comparison between two organisms.

Here, we take the next logical step to identify conserved (and divergent) transcriptional features of migratory NC cells between two organisms by directly comparing chick pharyngeal arches PA1 and PA2, isolated at HH13 with the front 20% and back 80% of the arches collected as distinct samples, with the externally protruding portions of mouse PA1 and PA2, isolated at E9.5. These tissue samples were analyzed jointly by scRNA-seq followed by canonical correlation analysis in Seurat (Stuart et al., 2019). Specifically, we performed an integrated analysis, determining the scRNA-seq clusters that emerged when the chick and mouse PA1-2 datasets were combined. We assigned identities to the clusters based upon expression of known marker genes and determined species-specific markers (chick only or mouse only) for each cluster. Cluster markers found in both the chick and mouse analyses were determined to be conserved between the two species. Our analyses uncovered conservation of Trailblazer signature genes within a small cluster containing both mouse and chick cells. We validate expression in the mouse of a subset of Trailblazer genes using multiplexed fluorescence in-situ hybridization and show gene knockdown disrupts NC cell migration into mouse PA1-2. Together, our comparative approach strengthens the theory that Trailblazer cells are an important feature of migratory NC cells and represents a roadmap to enhance and more deeply probe the NC cell GRN.

## RESULTS

### scRNA-seq and computational analyses successfully permit the direct comparison of chick and mouse pharyngeal arches

To compare NC cell gene expression profiles between the chick and mouse, we limited our analysis to genes that are clearly orthologous between the two species. In this process, we kept 72% of chick genes and 32% of mouse genes; a discrepancy likely accounted for by the greater read depth in the mouse experimental samples. When the chick and mouse datasets were combined utilizing Seurat integration, most clusters consisted of a clear mixture of cells from the two species (Fig. 1A,B; Table S1, S2) and cell numbers counted (Table S3). Clustering at 0.3 resolution resulted in 11 clusters, with identities assigned based on marker gene expression (Fig. 1C; Table 1). These data highlight the strengths of recent technological advances in single cell and computational analyses that now allow the direct comparison of chick and mouse pharyngeal arches.

**Fig 1.**
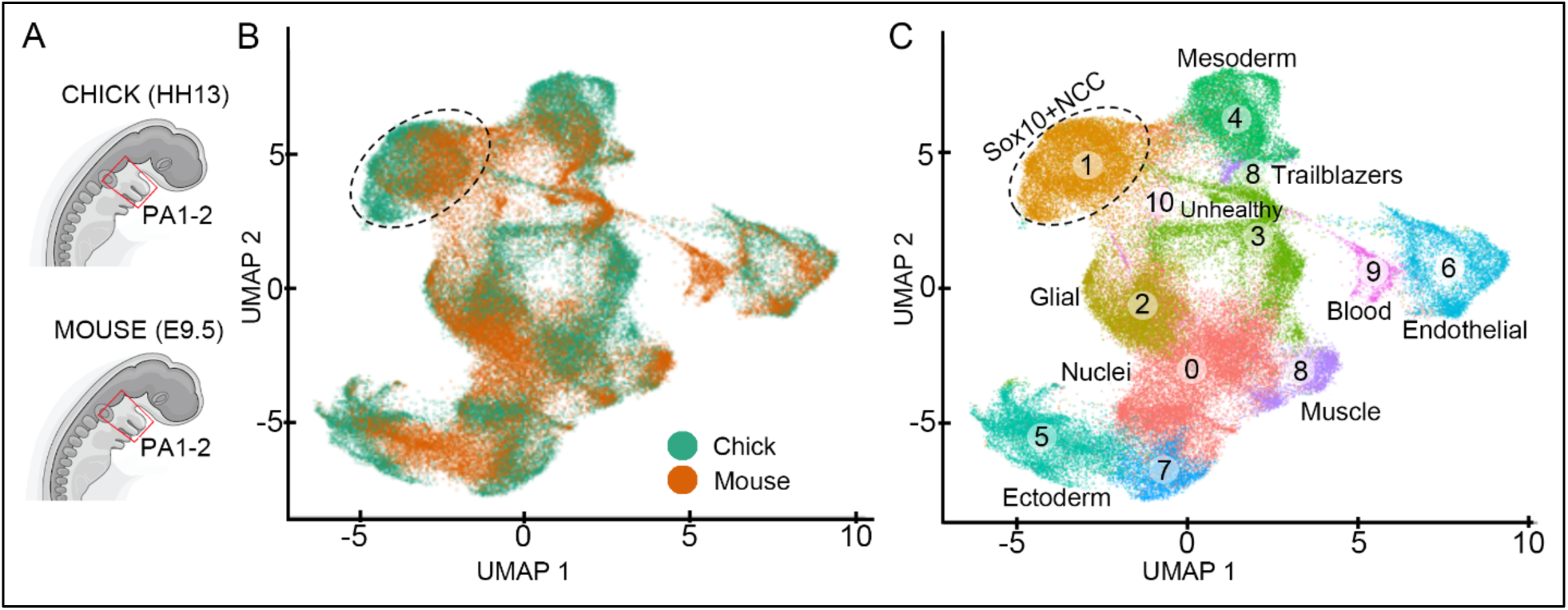
Clustering of mouse and chick datasets. (A) Schematic of tissue collection in the Chick (HH13) and Mouse (E9.5). (B) Joint clustering of mouse and chick datasets utilizing integrated analysis (in Seurat v3). (C) Cluster identity in integrated analysis of mouse and chick datasets. The oval dotted circle surrounds Cluster 1.

**Table 1.** Cluster identity based on marker gene expression in integrated mouse and chick analysis.

| Cluster | Identity | Marker Genes (Joint) | Marker Genes (Chick) | Marker Genes (Mouse) |
| --- | --- | --- | --- | --- |
| 0 | Nuclei | ID3, STMN1, H3F3B, HMGB2/3, UBB | ID3, STMN1, H3F3B, HMGB2/3, ERH, CALM2 | PRRX2, RMR2, MSX1/2 |
| 1 | Neural Crest | **LMO4**, **GJA1**, SOX10, ITGB3, TFAP2B | **LMO4**, SOX10, **GJA1**, ITGB3, CDH11, TFAP2B, COL9A1 | **LMO4**, **GJA1**, POLA2, BARX1, TWIST1, CRABP1 |
| 2 | Glial | FABP7, **CRABP-I**, DLX5, OAZ2, LDHB | FABP7/3, **CRABP-I**, OAZ2, LDHB | **CRABP-I**, DLX5, LDHA, FABP5 |
| 3 | Unhealthy Cells | **MT\***, **ND\***, CYTB, ATP6, COX3 | **MT\***, ATP6, **ND1/3/4/5/6, COX3** | **ND1/4/5/6**, CYTB, **MT\***, ATP6, **COX3** |
| 4 | Mesoderm | COL1A1, PITX2, EBF3, **FN1**, PRRX1, **VCAN** | COL1A1, PRRX1, EBF3/2, PITX2, **FN1**, **VCAN** | TSHZ2, **FN1**, LHX2, HLX, MSC, **VCAN** |
| 5 | Placodal Ectoderm | CLDN1/3, **EPCAM**, IGFBP2/5, FGF8, KRT18/24 | CLDN1/3, **EPCAM**, KRT24, ALCAM | IGFBP2/5, **EPCAM**, WNT6, KRT18, FGF8, CLDN7 |
| 6 | Endothelial | **LMO2**, **EGFL7**, **CDH5**, SOX18, **RAMP2** | CA2, **LMO2**, SOX18, **CDH5**, **RAMP2**, **EGFL7**, TAL1 | **RAMP2**, CLDN5, EMCN, VIM, **EGFL7**, **LMO2**, **CDH5** |
| 7 | Ectoderm | **KRT18/24/7**, **EPCAM** | **KRT18/24/7**, CMTM8, EZR, YBX3, **EPCAM** | **KRT18/24**, FOXE1, **EPCAM**, CLDN7, UCHL1 |
| 8 | Muscle | **ACTA2**, TNNI1, **GATA5**, **MYL9**, NKX2-5 | **ACTA2**, **GATA5**, NKX2-5, CSRP2, **MYL9** | **ACTA2**, ACTC1, **TNNI1**, TNNT2, **MYL9**, NKX2-5 |
| 8 | Trailblazer | BAMBI, KAZALD1, ISL1, DES | BAMBI, LUM, KAZALD1 | DES, ISL1 |
| 9 | Blood | HBE, HBA1, HBZ, **CA2**, **GBE** | **GBE**, **CA2**, TMSB4X | HBA1, HBE, HBZ, GBE, CA2, FECH |
| 10 | Neural Crest-derived Chondrocytes | BARX1, SOX9, COL1A2, COL2A1 | BARX1, COL1A2, POLA2, SOX9 |  |

**BOLD** genes are conserved cluster markers
NON-BOLD are species specific cluster markers

### Conserved cell types are validated within chick and mouse pharyngeal arches

We observed strongly defined clusters for endothelial cells, ectoderm, mesoderm, as well as NC cells and glia (Fig. 1B, C; Table 1). Aspects of our previously identified chick cranial NC cell invasion signature (Morrison et al., 2017; 2021) were present in Cluster 8, which is also characterized by expression of muscle markers (Fig. 1B, C; Table 1). Both chick and mouse cells were found in clusters that correspond to nuclei or unhealthy cells (Fig. 1B,C; Table 1, Cluster 3), which we excluded from further analysis. In contrast to the overall similarity, some clusters were unique to a specific organism. For example, we observed blood cells only in the mouse dataset (Fig. 1B,C; Table 1, Cluster 9) showing the specificity of the clustering methodology that recognizes blood cells as a mouse specific cluster. The majority of cells from the otic placode (Table 1, Cluster 5) were unique to the chick dataset. This was likely due to the inclusion of ‘proximal positioned’ cells in the chick NC cell migratory streams. Only distal regions of the mouse pharyngeal arches (PAs) were isolated, making the inclusion of any mouse otic placode cells less likely in these samples. These results validated our single cell analyses and encouraged more detailed investigation into the underlying GRN of neural crest development.

### Integrated analysis identifies conserved markers within the *SOX10*-positive NC cluster and mouse-specific markers, *NR2F1* and *BARX1* related to craniofacial development disorders

*SOX10* has been used as a canonical marker of migrating NC cells. However, we recently showed that *SOX10* expression decreases significantly as migrating chick NCs move farther away from the dorsal neural tube midline and into the pharyngeal arches (Morrison et al., 2021). In the mouse at E9.5, *SOX10* expression has a similar pattern; high expression in the proximal but not distal migrating NCs (Kuhlbrodt et al., 1998). This is consistent with the idea that by this stage in development some subpopulations of NCs have reached their future fate in the PAs. The presence of the ‘proximal’ portions of PA1 and PA2 in the chick data provide the majority of *SOX10* expression observed in these data. However, by looking at the expression of other NC cell markers such as *LMO4*, *TFAP2B*, and *TWIST1* in conjunction with *SOX10*, we successfully identify a cluster of NC cells containing both chick and mouse cells (Fig. 1C; Table 1).

Many of the top marker genes for the NC cell cluster (Fig. 1B, C; Table 1, Cluster 1) are genes that have previously been identified as NC cell markers in either the mouse or the chick, including *SOX10*, *LMO4*, *GJA1*, and *TWIST1* (Table 1). These genes consistently display bias to NC clusters in joint, chick only, and mouse only analyses (Table 1). The integrated analysis also identified NC marker genes such as *MYLK* and *OLFM1* which show NC-biased expression in the joint and chick clustering, but very limited expression in the mouse (Table S1), suggesting the possibility that these genes lack a conserved role in migratory NC cells.

The *SOX10*-positive NC cell cluster in the chick dataset was characterized by the expression of several marker genes that overlap with our previous work and analyses on the Trailblazer signature (Morrison et al., 2017, 2021) and the integrated analysis herein, including *SOX10*, *LMO4*, *GJA1*, and *CDH11* (Table 1). In contrast, other genes identified through this analysis appear to be characteristic of NCs in the chick, but are not statistically enriched in mouse NC cells, including *MYLK*, *ITGB3*, and *TNC* (Table S1).

Mouse genes enriched in NC cells (Cluster 1) include several canonical NC cell marker genes, including those that also typify NC cells in both the joint and chick analyses, such as *LMO4* and *GJA1* (Table 1). Additionally, this cluster is enriched for expression of genes which are not markers of chick NC cells, including *SOX9*, *POLA2*, *NR2F1*, and *BARX1* (Table S1). Interestingly, mouse NC cells are also enriched for *PRRX1* expression, which is present in the chick but more strongly expressed in the mesoderm than NC cells (Table S1). This is probably due to simple species-specific differentiation heterochrony.

To more rapidly visualize both species-specific and conserved markers of each cell type amid cluster definition, we created a dot plot that displays Table 1 markers across the scRNA-seq clusters (Fig. 2). We recognized markers that are shared between clusters. For example, CRABP1 is a marker of Cluster 1 (*SOX10*+NCs) and Cluster 2 (Glial) and *FN1* is a marker of Cluster 4 (Mesoderm) and Cluster 5 (Placodal ectoderm). In regard to Cluster 8 (Muscle/Trailblazer), *KRT18* and *VIM* are markers that are shared with Cluster 7 (Ectoderm) and Cluster 6 (Endothelial), respectively (Fig. 2). Further, *LMO2*, *CDH5, EGFL7* and *TMSB4X* are faithful markers of Cluster 6 (Endothelial) (Fig. 2) and markers of mouse only, chick only, and both species for each. In contrast, *ACTC1* and *TNNT2* mark muscle (Cluster 8) of mouse only but not chick (Fig. 2).

**Fig. 2.**
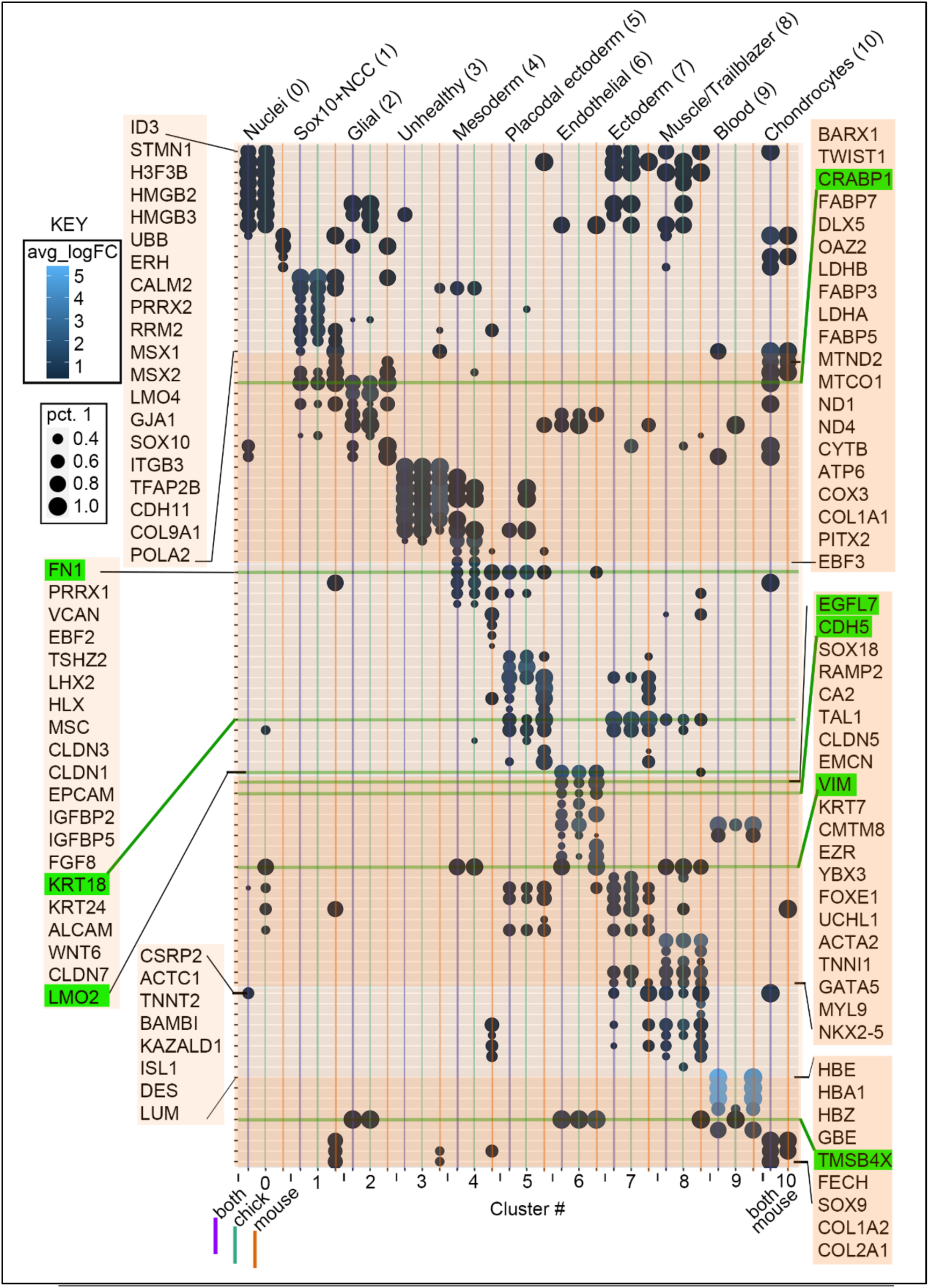
Dot plot displaying differential expression of Table 1 markers across all chick and mouse PA1-2 scRNA-seq clusters. (A) Each dot is colored by the average log fold change enrichment and sized by the percentage of cells expressing the gene. Species-specific as well as combined chick and mouse subpopulations of cells within each scRNA-seq cluster are shown, allowing rapid visualization of both species-specific and conserved markers of each cell type. The dot plot displays all Table 1 markers across joint and species-specific scRNA-seq clusters. For example, “both0” displays markers of chick and mouse cluster 0 cells, while ‘chick7’ represents markers of chick cluster 7 cells. The genes are ordered as in Table 1, which makes markers for each cluster visible along the diagonal. The cluster number is also in parentheses at the top.

### A NC cell invasion signature is conserved between chick and mouse PA1-2 migratory streams and includes enhanced expression of *KAZALD1, BAMBI, GPC3, and DES*

Previous gene profiling studies uncovered spatial molecular heterogeneities within the chick cranial PA2 NC cell migratory stream (McLennan et al., 2012; 2015). Single cell profiling of the NC cell migratory streams at progressive developmental stages, including the PA2 stream (Morrison et al., 2017) and all four PA1-4 streams (Morrison et al., 2021) then revealed a subpopulation of NCs within the migratory wavefront, termed trailblazers. Trailblazers expressed a transcriptional signature conserved throughout migration and included genes consistent with their role in cell invasion through the extracellular matrix and other cell types.

We sought to determine whether this invasion signature is conserved in migrating mouse cranial NC cells and considered the expression of the top 15 NC cell invasion genes across our integrated chick and mouse analyses (Fig. 1; Tables S1). We first noted that Cluster 8 cells appear in two, segregated locations on our UMAP and discovered cluster 8 markers were indicative of both muscle and Trailblazer NC cells (Fig. 3). Indeed, when we looked at the expression of individual cluster 8 markers, we observed chick PA2 trailblazer NC cell markers (based on Morrison et al., 2017), such as *(BAMBI*, *KAZALD1*, *DES*, *ISL1 and GPC3*) enriched in the small subset of cluster 8 attached to cluster 4 (Mesoderm) (Fig 3C, D; compared in B). When considering the majority of cluster 8 cells, we determined that muscle markers (*ACTA2* & *MYL9*), which are not chick PA2 trailblazer NC cell markers and not enriched in the small subset of cluster 8 attached to cluster 4 (Mesoderm), were enriched in the rest of cluster 8 (Fig. 3C, D; compared in B). Interestingly, this small cluster enriched for trailblazer markers identified in chick contains mostly mouse cells, highlighting conservation of trailblazer NC cells in both chick and mouse (Fig. 1B, C; Fig. 3). We confirm the expression of Gpc3, Bambi, Kazald1, and Des is restricted to leader mouse NCs in the 9.5 mouse mandibular arch by multiplexed FISH/HCR (Fig. 3E) and quantified by polyline kymograph analysis in the 5-slice projection (Fig. 3F, left) and whole image projection (Fig. 3F, right). This close association of transcriptional signatures between a subset of NCs and mesodermal cells recapitulates previous, independent findings in both chick and Zebrafish (Wagner et al., 2018; Morrison et al., 2021). This suggests that some biological properties are similar and shared between mesoderm and muscle with the neural crest.

**Figure 3.**
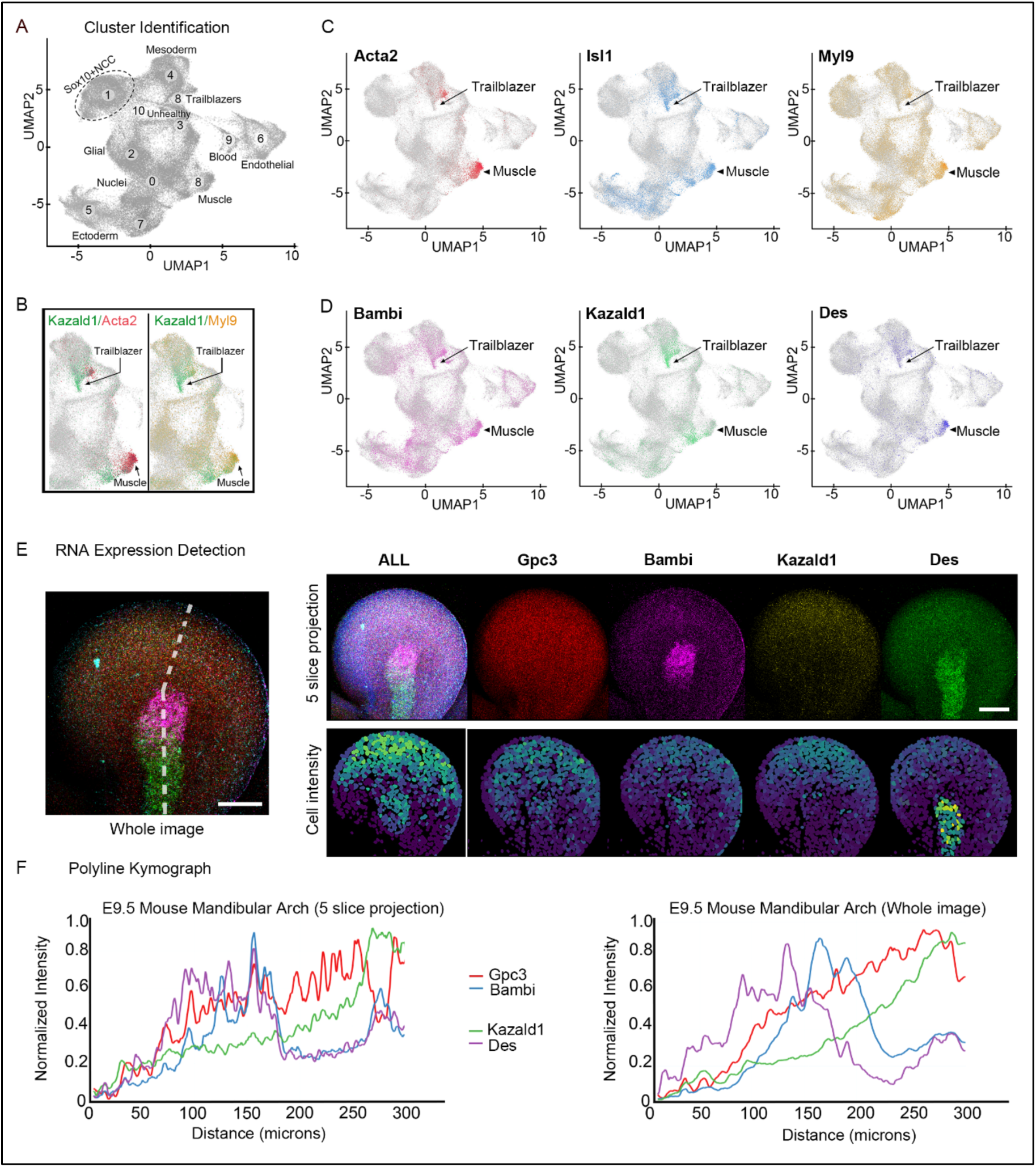
Feature plots that distinguish muscle and trailblazer NC cells in cluster 8 and subsequent HCR/polyline kymograph analysis in the E9.5 mouse mandibular Aach. (A) Cluster identification in grayscale as shown in Fig. 1A. (B) Overlay of two gene combinations selected from the individual images in (C-D) of all 6 genes plotted that are Top15 markers of cluster 8. Four out of the 6 genes are chick PA2 Trailblazer NC markers (based on Morrison et al., 2017) and enriched in the small subset of cluster 8 attached to cluster 4 (Mesoderm). Two out of the 6 genes are muscle markers (Acta2 & Myl9), which are not chick PA2 Trailblazer NC cell markers, and not enriched in the small subset of cluster 8 attached to cluster 4 (Mesoderm) but enriched in the rest of cluster 8. (E) E9.5 mouse mandibular arch labeled with HCR in situ for Gpc3, Bambi, Kazald1 and Des genes imaged with both channel and lifetime imaging. Left: maximum projection of all channels. Right: maximum projection of 5 slices from center of arch with DAPI segmentation colored by intensity of expression below. (F) Polyline kymographs reveal the proximal (left) to distal expression of genes highlighted in (E) along the line segment in (E). The dashed line is two segments to allow for the mandibular arch curvature where the line bends. The scalebars in E are 100 microns.

## SUMMARY

Examination of the expression of genes identified as part of the NC invasion signature across our analyses showed conservation of a cohort of genes, suggesting a common core invasion signature. This knowledge provides a foundation to study the functional roles of genes that drive NC cell invasion and migration into the pharyngeal arches. Interestingly, some of the genes identified from chick data also show strong expression and specificity for trailblazer lead NC cells in the mouse dataset including *KAZALD1* (Kazal-type serine peptidase inhibitor domain 1). *KAZALD1* encodes a secreted member of the insulin growth factor-binding protein (*IGFBP*) superfamily and it is deleted along with a small number of other genes in small chromosome 10q deletions associated with congenital anomalies, including cleft lip and palate – common neurocristopathies (Peltekova et al., 2014). The over-expression of *KAZALD1* has been correlated with high-grade glioma in human patient samples. Conversely lowering its levels in human glioblastoma cell lines (LL229 and U251) by treatment with siRNAs decreased cell invasion *in vitro* and *in vivo* in a nude mouse tumor xenograft model (Wang et al., 2013). Thus, *KAZALD1* may represent a novel molecular target to explore mechanisms regulating NC cell migration *in vivo*, the results of which may translate to a better understanding of human pathologies and cancer invasion.

We discovered many genes with expression domains that varied extensively across the two species; several of these genes showed expression in more than one cluster, most often in mesoderm and/or NC cells. This observation makes sense given that the NC cell invasion signature was initially identified in the chick data as clustering with mesodermal cells, which have recently been shown to move dynamically into the forming pharyngeal arches (McKinney et al., 2020). Single cell analysis has also highlighted the similarity of NC cells and mesodermal transcriptomes in Zebrafish (Wagner et al., 2018). Organism-specific expression of some NC cell invasion genes suggests there may be divergent processes and inputs between the mouse and chick, which will be a fascinating avenue for future investigation.

The joint clustering analysis identified several well-known marker genes but also revealed biases of NC cells in either the mouse or chick, suggesting this form of comparison may significantly enhance and influence future functional studies. For example, our integrated analysis identified *POLA2* and *NR2F1* as markers of mouse but not chick NC cells (Table 1; Table S1), suggesting they lack a conserved role in migrating NC cells. The role of *NR2F1* has been well-studied in human and mouse (Bergeron 2016; Rada-Iglesias et al. 2012), but not chick NC cells. Loss of *NR2F1* function results in abnormal migration of newly differentiated neurons, impacting cortical morphology and layer organization of the nascent cortical plate and is thought to underlie specific features of human neurodevelopmental disorders (Tocco et al., 2021). A second striking example in our joint clustering analysis was the assignment of *SOX9* as a mouse, but not chick NC cell marker (Table S1). A possible explanation for this difference in gene expression is that Meckel’s cartilage, the dentary bone, hyoid support structures and inner ear bones (or columella in the case of the chick) which are derived from first and second arch NC cells are most likely a reflection of specifies-specific differentiation heterochrony.

In summary, while the conclusions drawn here about the potential conserved and divergent roles of genes expressed during NC cell migration have provided new insights into NC biology and are consistent with previous observations about gene expression in the mouse and chick, they highlight the need for more comprehensive functional analysis to better understand the similarities and differences in the NC GRNs. Comparing the *in vivo* expression domains of genes of interest from this analysis will help to validate the biological significance of these observations. These data lay the foundation for future perturbation analyses and identification of downstream effects of knocking out a given gene, which can experimentally probe the conservation (or divergence) of the functional role of the gene at this stage of NC cell development. A combinatorial approach focusing on assaying key genes of interest in this way across model organisms will ultimately lead to a more detailed elaboration of GRNs for NC cell development across different model organisms and yield insights into the conservation of the processes involved. Together, this knowledge has the potential to inform studies of human neural crest cell migration-related birth defects and strategies for devising stem-cell based tissue repair.

## MATERIALS AND METHODS

### Single cell isolation of chick tissue

Four chicken samples and two mouse samples of 10x were sequenced on a total of 9 Illumina HiSeq 2500 flow cells (Illumina, San Diego, CA, USA). Specifically, fertilized, white leghorn chicken eggs (NCBI Taxonomy ID:9031; Centurion Poultry, Lexington, GA, USA) were incubated at 38 degrees C in a humidified incubator until the desired Hamburger and Hamilton (HH) stage (Hamburger and Hamilton, 1951). Healthy embryos were harvested into chilled 0.1% DEPC phosphate-buffered saline (PBS). To capture neural crest during active migration, pharyngeal arches (PA) 1 and 2 were manually isolated from HH13 embryos (n=15 embryos). Stage-, pharyngeal arch- and tissue portion-matched tissues were pooled and dissociated as previously described (Morrison et al., 2017). The viability and concentration of each single cell suspension was quickly confirmed on a Nexelome Cellometer Auto T4 (Nexelome Bioscience, Lawrence, MA, USA) and the cell suspensions used as input for 10x scRNA-seq (10x Genomics, San Francisco, CA, USA) per the manufacturer’s recommendations.

### Single cell isolation of mouse tissue

All animals were maintained by Laboratory Animal Services (LAS) at Stowers Institute for Medical Research (SIMR) and all experiments involving mice were approved by the Institutional Animal Care and Use Committee (IACUC) of the Stowers Institute for Medical Research under a protocol (Protocol 2021-134) issued to R.K. Euthanasia procedures were performed in accordance with recommendations by the American Veterinary Medical Association. To reduce variability due to differences in developmental stages, embryos selected for analysis and dissection were determined to be E9.5 only if they had between 22-24 somite pairs.

Pharygeal arches 1 and 2 from wildtype E9.5 CBA/Ca/J x C57BL/10 mouse embryos were manually dissected in ice-cold DEPC-treated phosphate-buffered saline (DPBS, Sigma catalog number D8537). Individual PAs were pooled into two samples (PA1 and PA2), which were dissociated by incubation in 0.25% Trypsin + EDTA (Gibco catalog number 25200-056) for 3 minutes with two rounds of manual disruption by pipetting. The reaction was stopped with fetal bovine serum and samples were washed two times in DPBS. Single-cell suspensions were loaded on a Nexelome Cellometer Auto T4 to assess cell count and viability, then used as input for 10x scRNA-seq.

### 10x Chromium single-cell RNA-seq library construction

Dissociated cells were loaded on a Chromium Single Cell Controller (10x Genomics, Pleasanton, CA, USA), based on live cell concentration, with a target of 4,000-10,000 cells per sample. Libraries were prepared using the Chromium Single Cell 3’ Library & Gel Bead Kit v2 (10x Genomics, Pleasanton, CA, USA) according to manufacturer’s directions. Resulting short fragment libraries were checked for quality and quantity using an Bioanalyzer 2100 (Agilent Technologies, Santa Clara, CA, USA) and Qubit Fluorometer (Invitrogen, Waltham, MA, USA). Libraries were pooled and sequenced to a depth necessary to achieve 25-35,000 mean reads per cell, 125-520M reads each, on an Illumina HiSeq 2500 (Illumina, San Diego, CA, USA) instrument using Rapid SBS v2 chemistry with the following paired read lengths: 26 bp Read1, 8 bp I7 Index and 98 bp Read2.

### Bioinformatics

Data was processed with bcl2fastq2 (2.20) and aligned with CellRanger (3.0.0). Chicken data was aligned to galGal6 from UCSC using annotations from Ensembl 98. Mouse data was aligned to mm10 from UCSC using annotations from Ensembl 98. Downstream analysis was done in R (3.6.1) with the Seurat package (3.1.5.9003) (Butler 2018). Cells were kept for downstream analysis if they had more than 500 genes expressed, less than 3k genes expressed for chicken, less then 6k genes expressed for mouse, and less then 20% mitochondrial expression. Data was analyzed using the sctransform function in Seurat, and the first 40 principal components were used for later steps (UMAP and cluster identification).

To compare chicken and mouse orthologous genes, we downloaded orthologous gene information from Ensembl (98) BioMart. In cases where multiple orthologous genes were identified between chicken and mouse, the first mouse gene was kept based on percent identity with the chicken gene. In total, we had 13,524 genes in our orthologous gene table. Single cell data for genes that did not have orthologous gene matches was discarded for this analysis.

After individual analysis, chicken and mouse datasets were integrated using the Seurat IntegrateData function. Clusters were identified using Seurat FindClusters with resolution of .5. Marker genes for clusters were identified using Seurat FindAllMarkers with only.pos set to TRUE to find positive markers, and the minimum percentage set to 0.25 and log fold change threshold set to 0.25.

Application of several data integration analysis tools failed to generate consistent results. We proceeded with Seurat’s CCA, which is commonly used in the field and yielded the best integration of our data sets.

### HCR and Imaging

Tissue was dissected and fixed in 4% PFA at the appropriate stages. Hybridized Chain Reaction HCR was performed according to Molecular Instruments protocol for GPC3 (ENSMUSG00000055653), Kazald1 (ENSMUSG00000025213), Bambi (ENSMUSG00000024232) and Des (ENSMUSG00000026208) using probes designed to amplifiers B3, B2, B1 and B4 respectively and labeled with AlexaFluor 647, 594, 555 and 546 respectively. Images were acquired using a combination of channel and lifetime imaging on a Leica SP8 confocal microscope equipped with an HC PL APO CS2 40x objective lens and lifetime rig. Channels labeled with AlexaFluor 555 and 546 were separated with fluorescence lifetime and remaining channels, including DAPI, were imaged by channels. Full z-stacks at 2um step size were taken of 4 mandibular and 4 hyoid arches. Polyline kymographs were performed in Fiji on either the center 5 slices of the image or full z-stack projections. Cell segmentation and intensity analysis was performed in python using Cellpose.

## Acknowledgements

The authors thank Christof Nolte and Paul Trainor for helpful discussions and feedback on the project and manuscript; Heidi Monnin, Haley Tansey and Ian Castillo-Jolly for mouse husbandry. We especially thank the Microscopy, Imaging and Big Data, Laboratory Animal Services, and Sequencing and Discovery Genomics technology centers at the Stowers Institute for their help in supporting this project. This work was conducted to fulfill, in part, the requirements for I.P.’s degree as a PhD student with the the Graduate School of the Stowers Institute. This work was supported by funds from the Stowers Institute for Medical Research to R.K. (grant number 1001) and to P.M.K., and partial support to P.M.K. (NIH R21 HD106033). Original data underlying this manuscript can be accessed from the Stowers Original Data Repository at [http://odr.stowers.org/websimr/].

---

**Supplemental Table S1:** Top 15 markers for each of the 11 cluster in the UMAP.

**Supplemental Table S2:** Full marker list for integrated analysis from 11 cluster UMAP. (See attached file).

**Supplemental Table S3:** Cell counts for the profiling analyses.

**Table S1.**
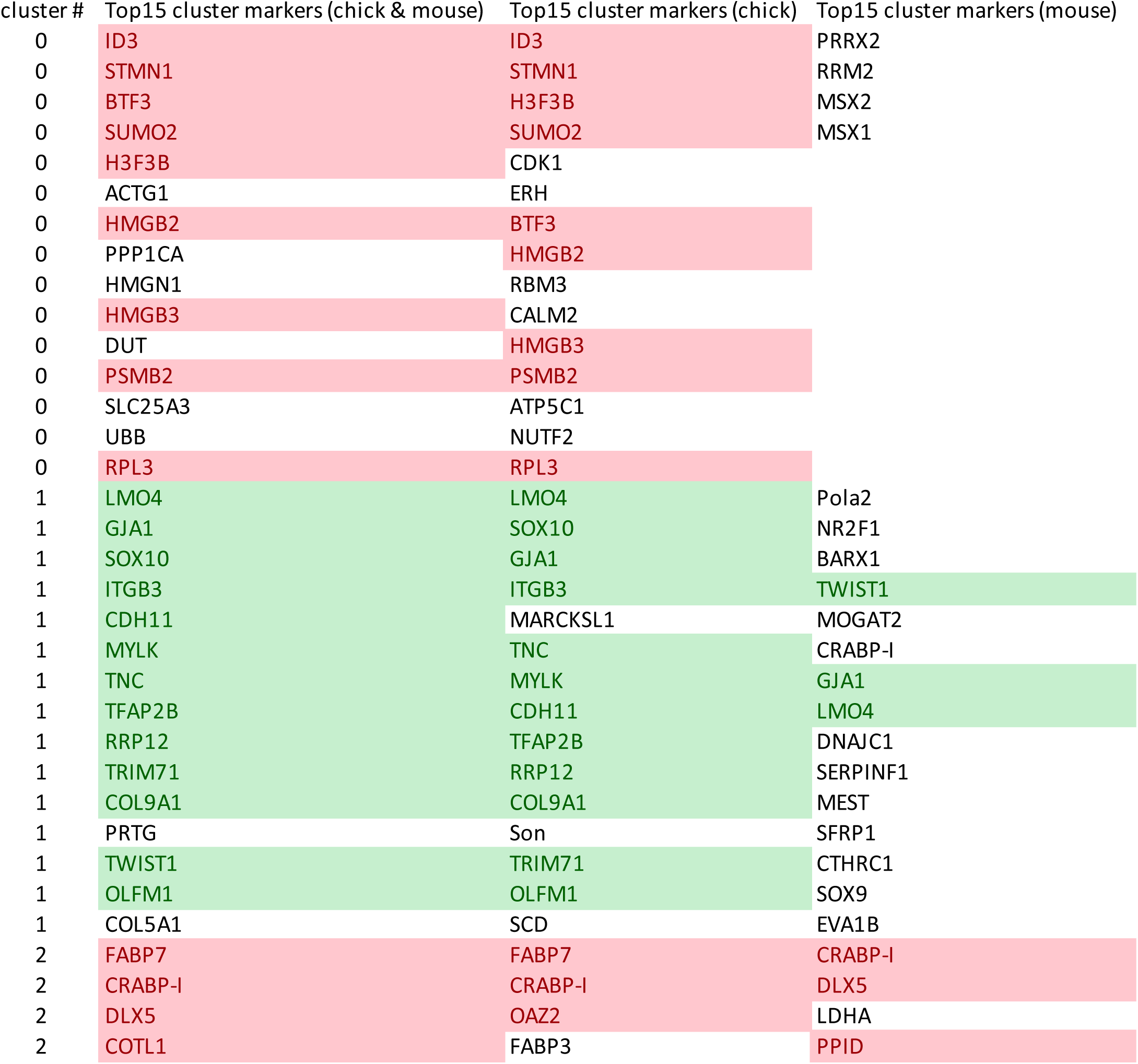
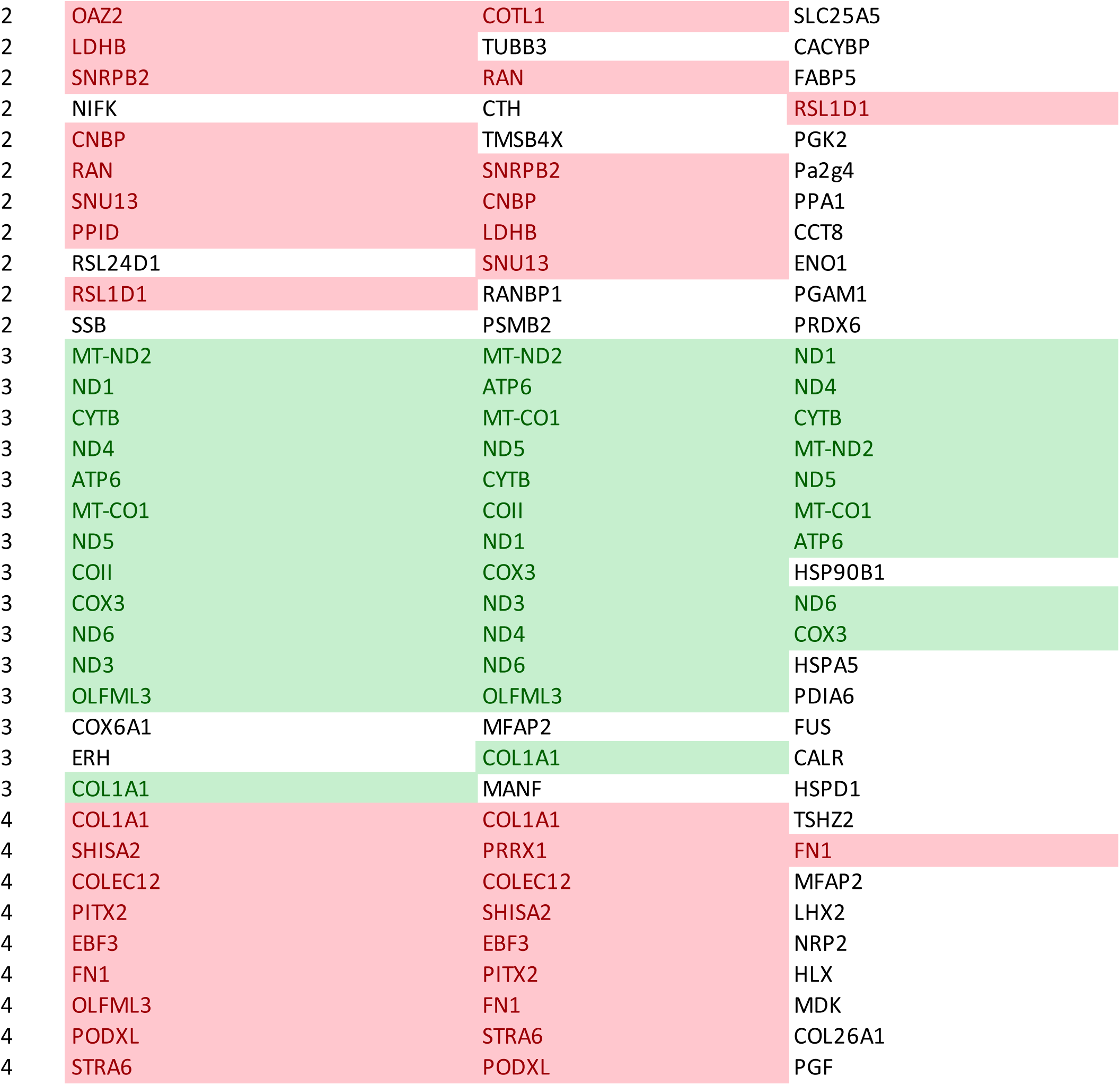
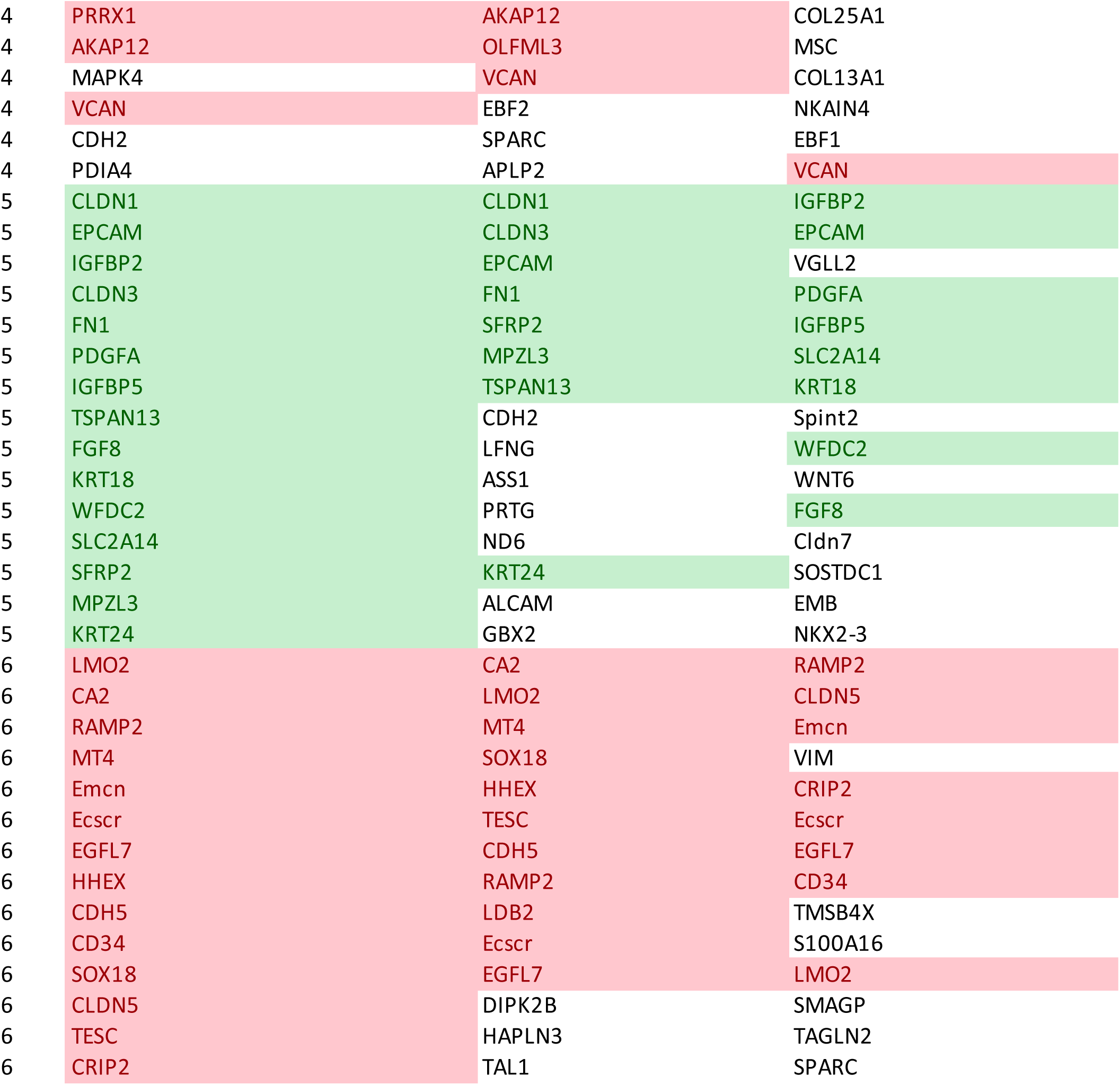
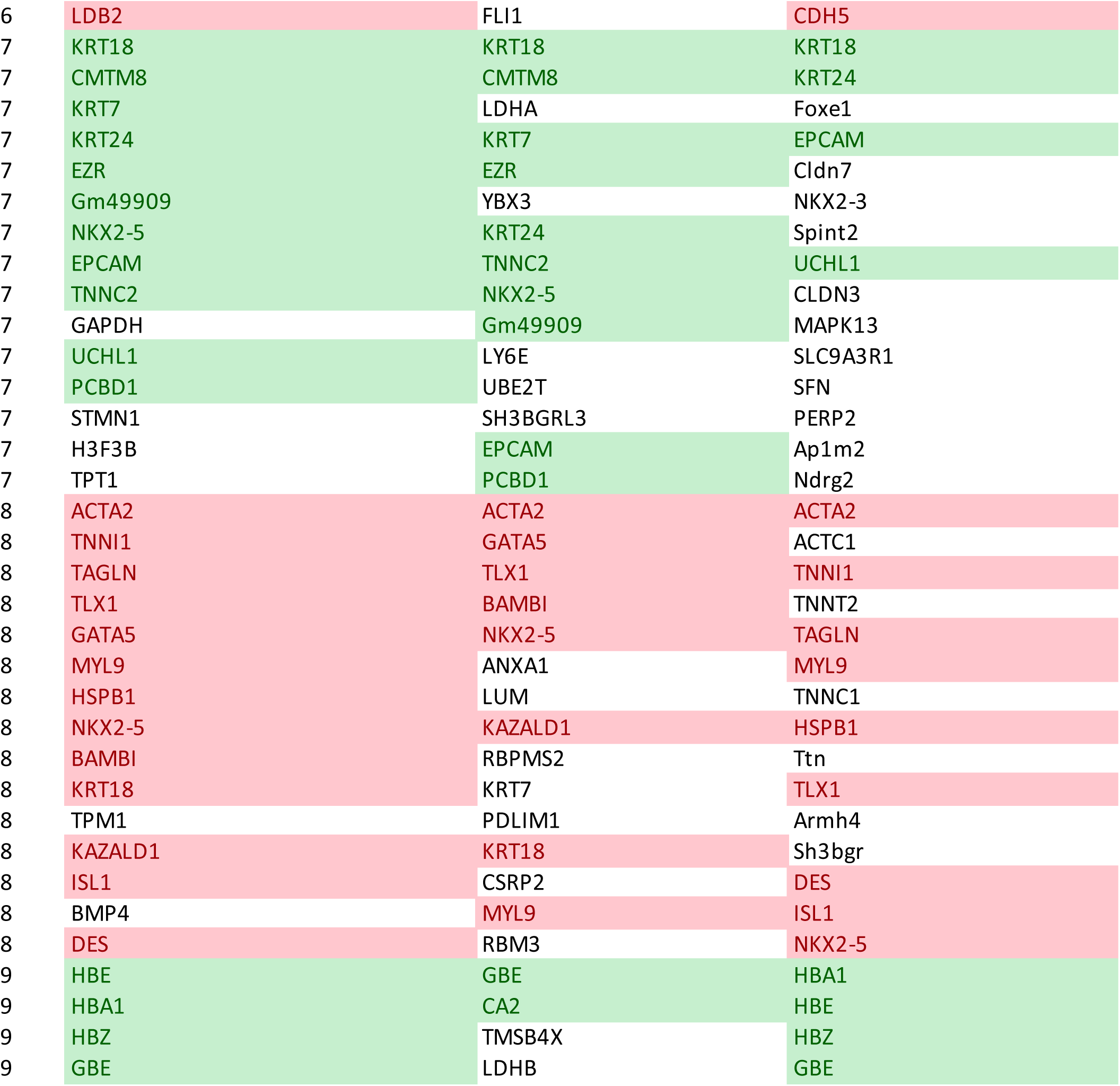
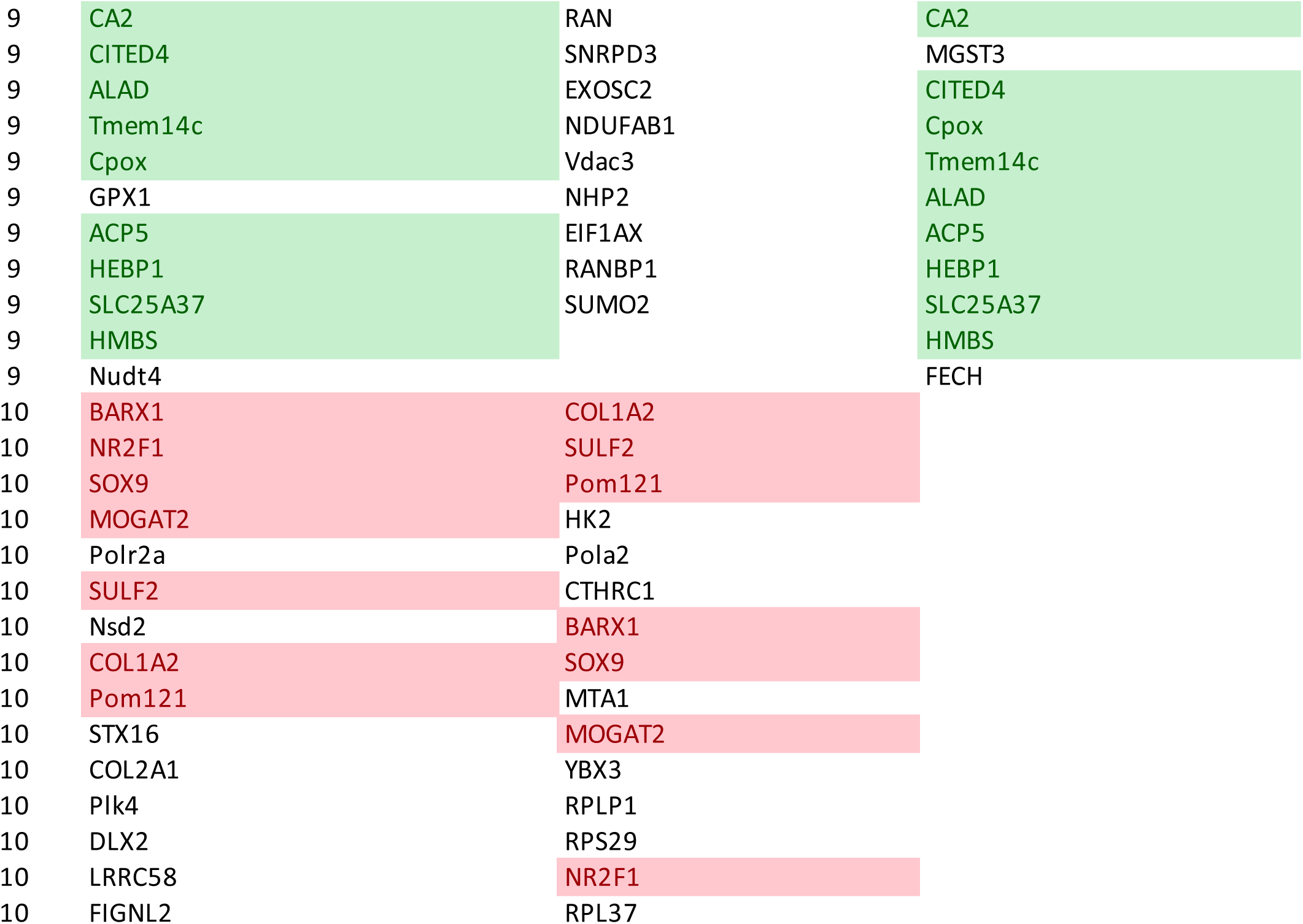
Highlighted genes overlap between Joint (chick & mouse) and at least one species for that cluster.

**Table S3:**
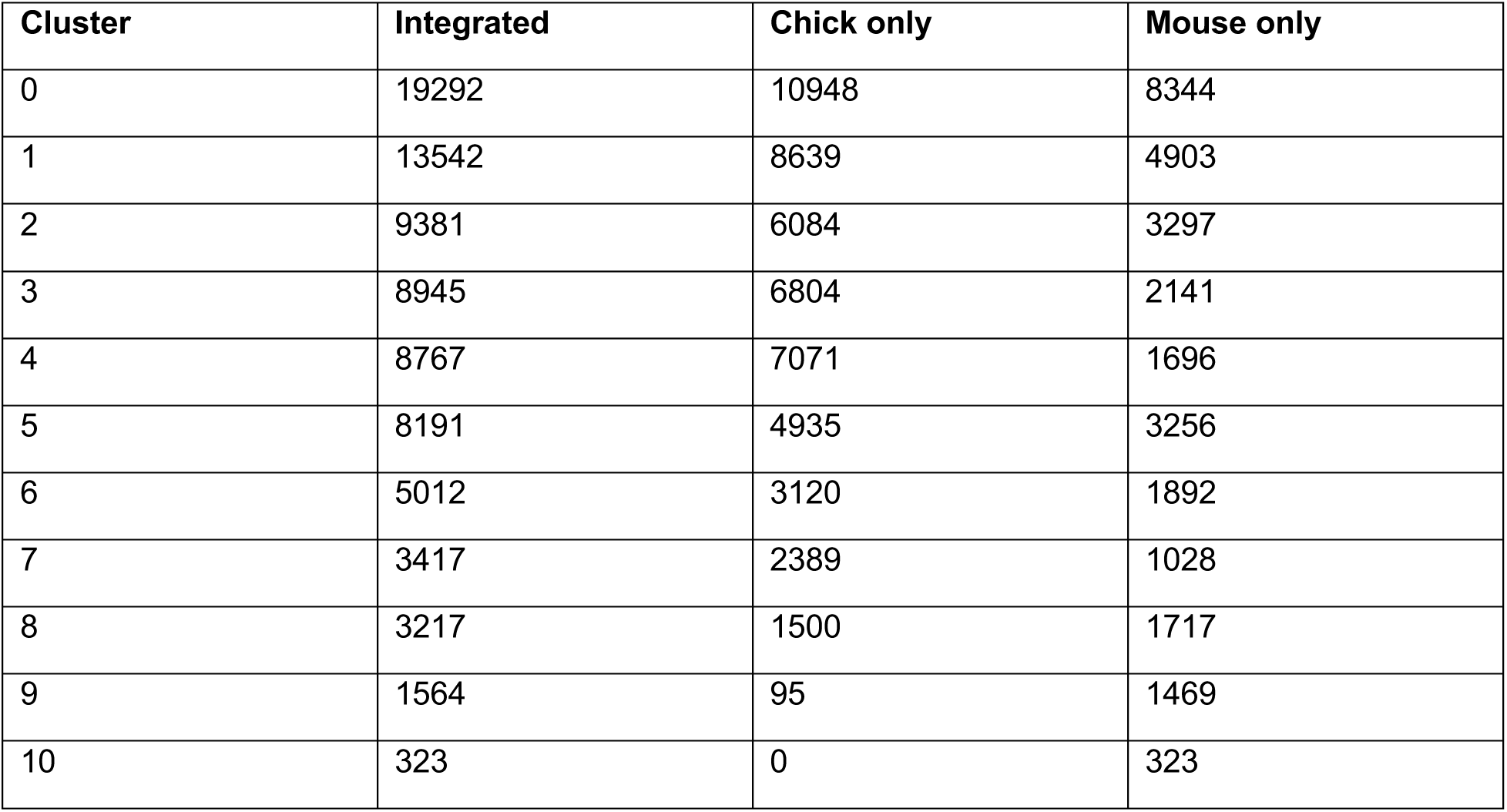
Cell counts for the profiling analyses.

